# Orbitofrontal Cortex Modulates Auditory Cortical Sensitivity and Sound Perception

**DOI:** 10.1101/2023.12.18.570797

**Authors:** Matheus Macedo-Lima, Lashaka Sierra Hamlette, Melissa L. Caras

## Abstract

Sensory perception is dynamic, quickly adapting to sudden shifts in environmental or behavioral context. Though decades of work have established that these dynamics are mediated by rapid fluctuations in sensory cortical activity, we have a limited understanding of the brain regions and pathways that orchestrate these changes. Neurons in the orbitofrontal cortex (OFC) encode contextual information, and recent data suggest that some of these signals are transmitted to sensory cortices. Whether and how these signals shape sensory encoding and perceptual sensitivity remains uncertain. Here, we asked whether the OFC mediates context-dependent changes in auditory cortical sensitivity and sound perception by monitoring and manipulating OFC activity in freely moving animals under two behavioral contexts: passive sound exposure and engagement in an amplitude modulation (AM) detection task. We found that the majority of OFC neurons, including the specific subset that innervate the auditory cortex, were strongly modulated by task engagement. Pharmacological inactivation of the OFC prevented rapid context-dependent changes in auditory cortical firing, and significantly impaired behavioral AM detection. Our findings suggest that contextual information from the OFC mediates rapid plasticity in the auditory cortex and facilitates the perception of behaviorally relevant sounds.

**Significance Statement:** Sensory perception depends on the context in which stimuli are presented. For example, perception is enhanced when stimuli are informative, such as when they are important to solve a task. Perceptual enhancements result from an increase in the sensitivity of sensory cortical neurons; however, we do not fully understand how such changes are initiated in the brain. Here, we tested the role of the orbitofrontal cortex (OFC) in controlling auditory cortical sensitivity and sound perception. We found that OFC neurons change their activity when animals perform a sound detection task. Inactivating OFC impairs sound detection and prevents task-dependent increases in auditory cortical sensitivity. Our findings suggest that the OFC controls contextual modulations of the auditory cortex and sound perception.

## Introduction

Sensory perception is highly flexible, constantly adjusting to context-dependent changes in arousal (1–4), attention (5, 6), or expectations about upcoming events (7–12). This rapid flexibility favors the detection of important or informative stimuli, supporting improved communication, predator avoidance, and reproductive success.

Context-dependent shifts in perception are thought to result from rapid changes in sensory cortical processing. For instance, when a stimulus suddenly acquires behavioral relevance (such as when it must be used to solve a task), sensory cortical neurons quickly adjust their tuning properties, response strengths, and/or functional connectivity to optimize its detection, discrimination, or identification (6, 13–31). These data indicate that sensory cortices must receive signals that tell them when and how to adjust their activity, but where and how such signals are initiated remains unclear.

Several lines of evidence suggest that the orbitofrontal cortex (OFC) is particularly well-positioned to mediate contextual signaling. First, OFC is unique in that it is the only frontal cortical region to integrate inputs from all sensory modalities (32, 33), providing it a distinct vantage point for discerning shifts in environmental or behavioral demands. Second, in addition to their well-established role in encoding value (34), OFC neurons also encode social (35), spatial (36, 37), and behavioral context (38–41). While recent work indicates that some of these signals are transmitted to sensory cortices (38–41), their downstream impact on sensory encoding and perceptual sensitivity is only beginning to be understood.

Here, we asked whether OFC shapes sound processing and perception by relaying contextual information to the auditory cortex in freely moving animals. We first recorded extracellular activity from OFC neurons under two different behavioral contexts: passive sound exposure and engagement in a sound detection task. We found that the majority of OFC neurons, including the specific subset that innervate the auditory cortex, exhibited context-dependent changes in firing rates. We then pharmacologically suppressed OFC activity during task engagement and assessed the effect on auditory perception and auditory cortical sound processing. OFC inactivation both impaired behavioral sound detection and prevented task-dependent changes of auditory cortical firing. Our findings suggest that the OFC supports context-dependent modulations of stimulus representations within the auditory cortex, contributing to the enhanced perception of behaviorally relevant sounds.

## Results

### OFC neurons are sensitive to behavioral context

Auditory cortical neurons are more sensitive to amplitude-modulated (AM) stimuli when animals perform an AM detection task compared to when the same sounds are presented in a passive, non-task context (13). This finding, which is consistent with decades of work (17, 18, 20, 22, 23, 27, 28, 42–46), suggests that task engagement recruits non-sensory networks that modulate auditory cortical response properties. If OFC participates in this process, then we should also observe a neural signature of task engagement in OFC neurons.

To test this prediction, we trained Mongolian gerbils on the same AM detection task described above. In this task, animals must drink from a waterspout in the presence of unmodulated noise and withdraw from the spout when the sound transitions to an AM noise to avoid an aversive shock (Figure 1A). Animals learned the task quickly, reaching expert performance levels (d’ *≥* 2, see Materials and Methods) within ∼250 trials across six sessions (Figure 1B-C; n = 16 animals across all experiments). We then implanted five of these animals (two female) with 64-channel silicon probe arrays in left OFC and recorded from 298 single- and 227 multi-units during task engagement and during passive sound exposure sessions just before (‘passive-pre’) and just after (‘passive-post’) the task (Figure 2A-C). During passive exposure sessions, the spout was removed from the test arena and the shocker was turned off, but everything else, including the sound stimuli and position of the recording electrodes, remained identical to the task.

**Figure 1.**
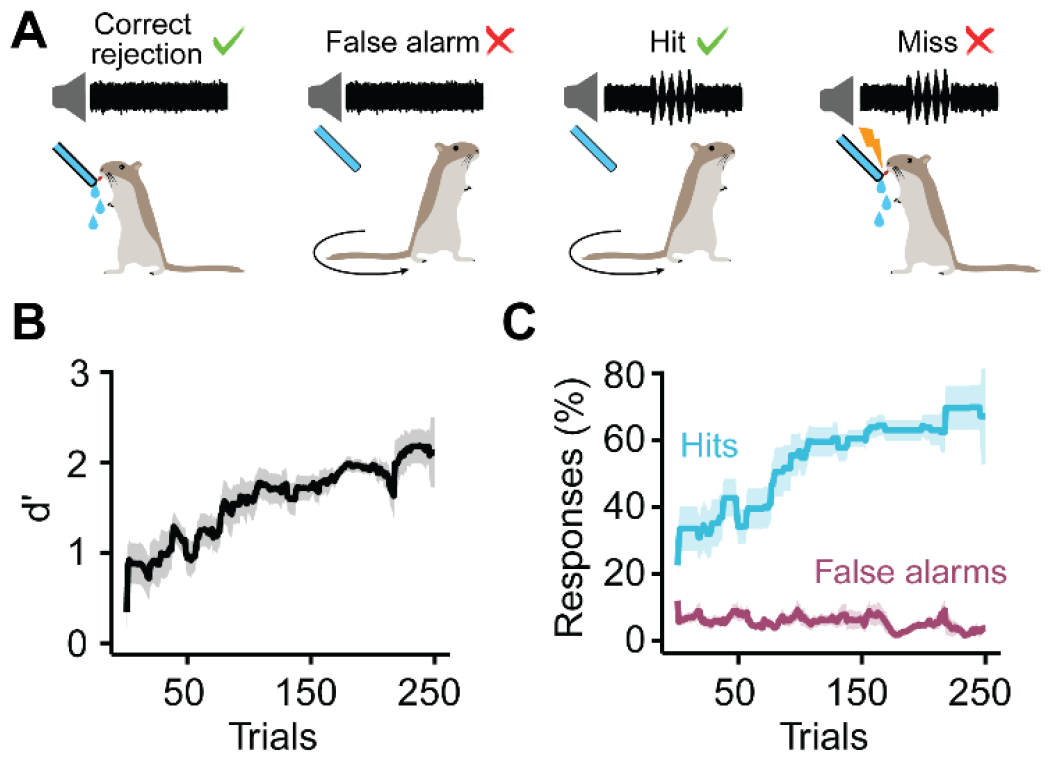
Behavioral task. (A) Task structure. Water-deprived animals were trained to drink from a waterspout while unmodulated noise continuously played from a speaker, and to withdraw from the spout upon presentation of amplitude-modulated (AM) noise. Failing to withdraw during AM noise was punished with a mild shock. (B-C) Mean ± standard error *d’* values (B) or hit rates and false alarm rates (C) for all animals included in this study (n = 16). Data were concatenated from training sessions over six days and analyzed using a 20-trial sliding window. Animals were only used for experiments after they reached expert performance levels (d’ *≥* 2).

**Figure 2.**
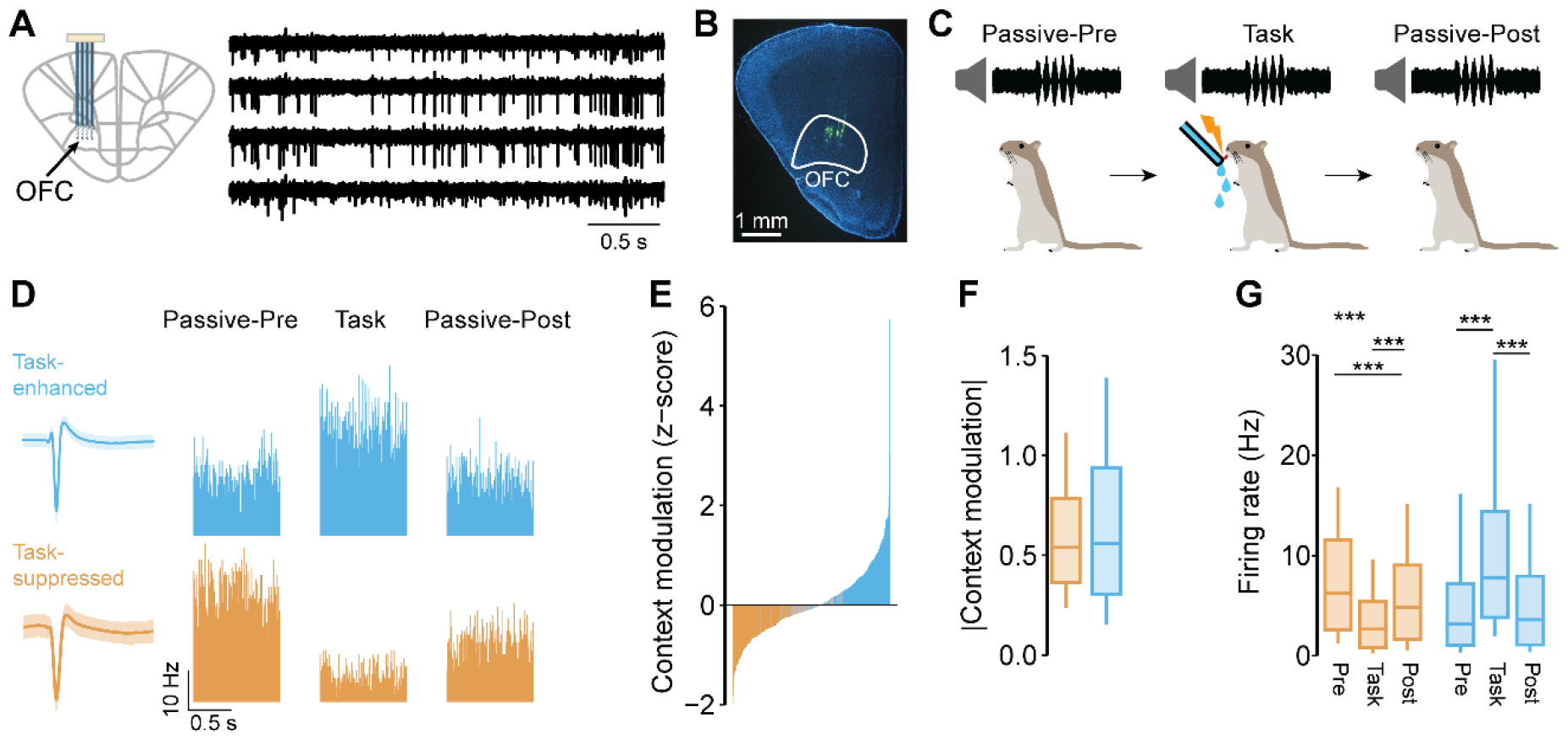
OFC neurons are sensitive to behavioral context. (A) Extracellular recordings were made from 64-channel electrode arrays chronically inserted into the left OFC. Sample recordings illustrate filtered voltage traces from four adjacent channels capturing activity from the same unit. (B) Histological images were used to confirm electrode placement by visualizing fluorescently labeled tracks (green) in the OFC. (C) OFC activity was recorded during task engagement and during passive sound exposure sessions just before (‘passive-pre’) and just after (‘passive-post’) the task. The same units were recorded across all three sessions. (D) Peristimulus time-histograms of two representative OFC neurons illustrate the effect of task engagement on non-AM firing rates. The ‘task-enhanced’ neuron (blue) exhibited a significant increase in firing during the task, while the ‘task-suppressed’ neuron (orange) exhibited a significant decrease. Mean ± standard deviation waveforms for each unit are shown on the left. (E) The magnitude of context modulation for each unit was calculated as the difference between the non-AM firing rate during the task and the non-AM firing rate during the passive-pre session z-scored over trials. Modulation values are presented for all units sorted in increasing order. Gray bars represent units without significant contextual modulation. (F) Absolute modulation magnitudes were similar between task-enhanced and - suppressed units. (G) Task-suppressed units exhibited reductions in non-AM firing that persisted into the passive-post period. The activity of task-enhanced units, on the other hand, returned to passive-pre levels after the task ended. ***p<0.001

We first examined the effect of task engagement on OFC activity during the presentation of AM stimuli. During passive exposure sessions, OFC firing was unremarkable, with no obvious change evoked by AM sounds. During task engagement, on the other hand, OFC neurons exhibited a heterogenous array of responses (Figure S1). While some of these neurons may have been sensitive to the AM stimulus, the majority appeared to be more sensitive to behavioral outcome, with changes in firing sometimes lasting 2-4 seconds after AM onset. To verify whether the OFC neurons that innervate auditory cortex exhibit similar dynamics, we injected a virus into the left auditory cortex to retrogradely express the calcium indicator jGCaMP8s and implanted an optical fiber into the left OFC (n = 9; four females). We found that during passive sound exposure, calcium transients were largely absent. However, during task engagement, calcium transients were robust and sensitive to behavioral outcome (Figure S2). These findings, which are broadly consistent with prior work (38, 40, 41, 47) confirm that OFC neurons, including the subset of OFC neurons that project to auditory cortex, encode behaviorally relevant information during task engagement.

We next examined the effect of behavioral context on the firing rates of OFC units during periods of unmodulated noise presentation. We found that most units recorded (397/525; 76%) were significantly modulated by context. Overall, 200/525 units (38%) were ‘task-enhanced’, exhibiting elevated firing rates during the task compared to the initial passive-pre session (Figure 2D; Wilcoxon rank-sum test for each unit; see Supplementary Methods in SI Appendix). A roughly equal number (197/525; 38%) were ‘task-suppressed’. A smooth continuum of context-dependent modulation was clearly observed (Figure 2E), similar to previous reports in the auditory cortex (48). Moreover, the magnitude of the context modulation was largely similar between task-enhanced and -suppressed units (p = 0.207; Figure 2F).

The effect of task engagement on auditory cortical neurons can persist for several minutes or hours following task termination (13, 23–25, 30, 49). If OFC mediates the changes observed in auditory cortex, OFC neurons should exhibit similar temporal dynamics. Therefore, we asked whether changes in OFC activity persist into the passive sound exposure session immediately following the task. As shown in Figure 2G, the firing rates of task-enhanced units returned to passive-pre levels after the task was over (p > 0.999, see Table 1 for full statistical results). In contrast, task-suppressed units exhibited longer-lasting changes, such that firing rates during the passive-post session remained significantly lower than during the passive-pre session (p < 0.001). All findings above were replicated when our analyses were restricted to single-units (Figure S3).

**Table 1.**
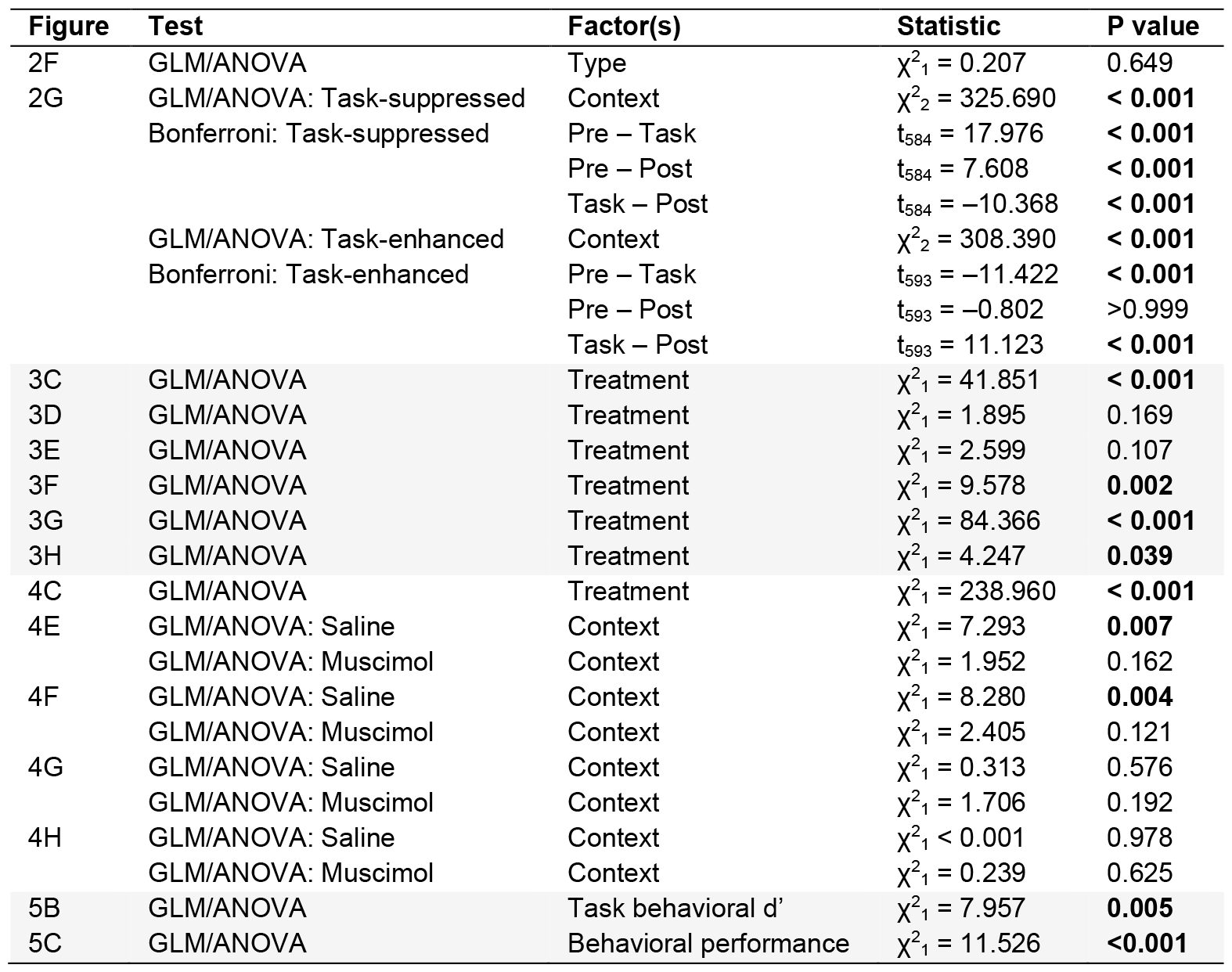
Statistical results.

Together, these data reveal that a subset of OFC neurons exhibit task-dependent modulations that evolve over a similar time course as those observed in the auditory cortex and raise the possibility that task-dependent changes in OFC and the auditory cortex are mechanistically linked.

### OFC inactivation impairs AM detection behavior

Task-dependent modulation of the auditory cortex is believed to enhance the detection or discrimination of behaviorally relevant stimuli (13, 17, 18, 20, 22, 23, 27, 28, 42–46, 48). If OFC mediates task-dependent modulations in the auditory cortex, then suppressing OFC activity should impair sound detection. To test this prediction, we implanted cannulas in bilateral OFC of nine gerbils (six females) previously trained to detect AM stimuli. In one male and one female, right hemisphere infusion cannulas clogged, but data from these animals were still included to evaluate the effect of unilateral left OFC inactivation. On alternating days, we infused either saline or muscimol (a GABA_a_ agonist that reversibly silences neural activity) into the OFC shortly before animals performed the AM detection task (Figure 3A-B).

**Figure 3.**
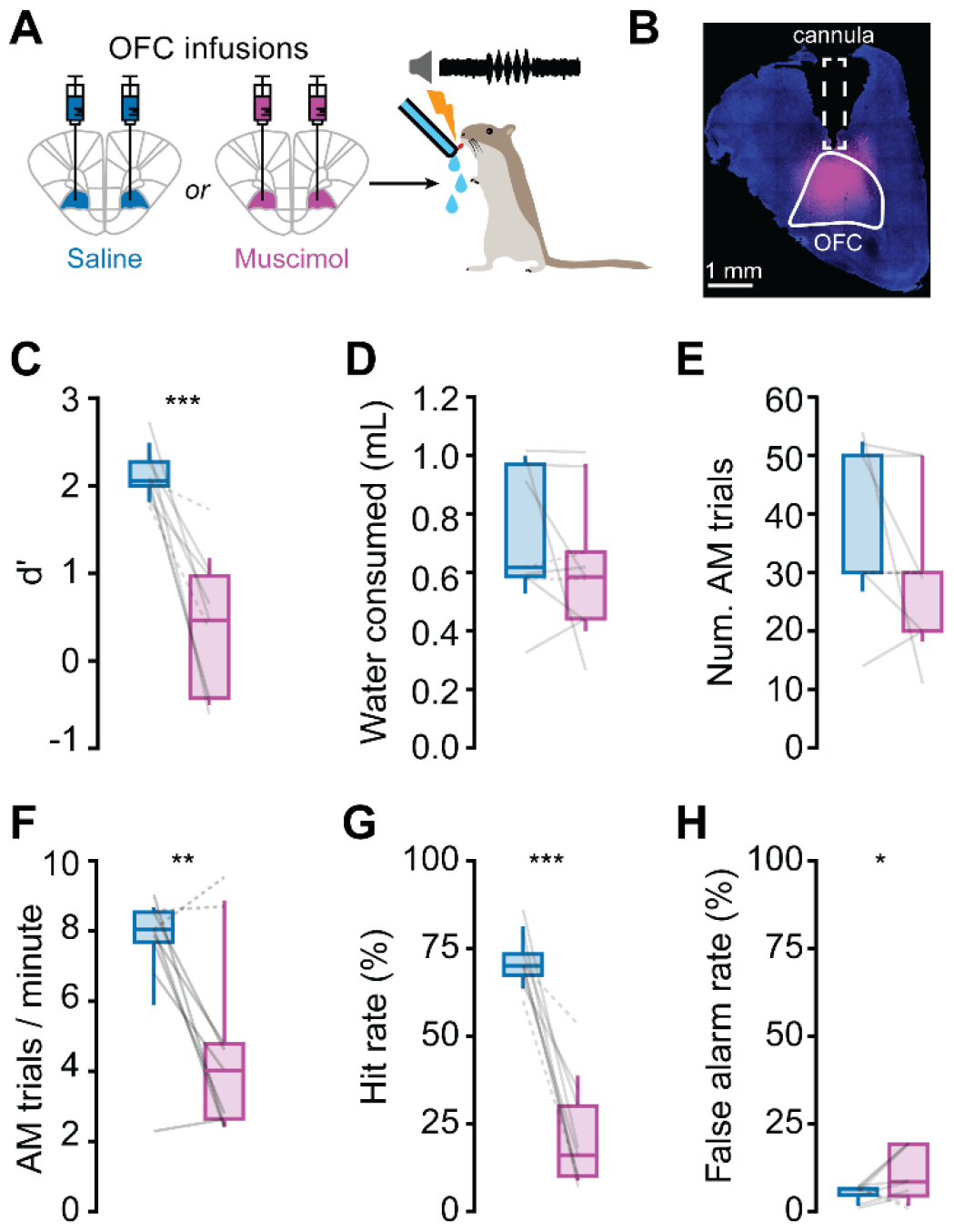
OFC inactivation impairs behavioral AM detection. (A) On alternating days, muscimol or saline was infused into bilateral OFC ∼40 minutes before animals performed the AM detection task. (B) At the conclusion of the experiment, a fluororuby infusion just prior to transcardial perfusion provided an estimation of drug spread. (C-E) Muscimol infusions reduced behavioral *d’* values (C), without systematically affecting water consumption (D) or the number of completed AM trials (E). However, muscimol infusion did reduce the rate of AM trial completion (F). (G-H) Muscimol infusions significantly reduced hit rates (G), and only slightly increased false alarm rates (H). Dashed lines indicate data from animals that received unilateral left infusions due to partially clogged cannulas. *p<0.05, **p<0.01, ***p<0.001.

OFC inactivation significantly impaired AM detection. As shown in Figure 3C, behavioral *d’* values were lower (poorer) in all animals tested after muscimol was infused into the OFC, compared to when saline was infused (p < 0.001; see Table 1 for full statistical results). Impaired performance could not be explained by reduced motivation to engage in the task, as the amount of water consumed (Figure 3D) and the number of AM trials completed (Figure 3E) were unaffected by muscimol infusion (both p > 0.1). However, OFC inactivation did reduce the rate of trial completion (p = 0.002, Figure 3F). This observation might be explained by the fact that the poor performance caused by muscimol infusion resulted in animals receiving more frequent punishment, thereby generating more frequent (and longer) spout withdrawals. Indeed, miss rates strongly predicted rates of trial completion (p < 0.001; Figure S4A).

OFC inactivation may have impaired behavioral performance by disrupting an animal’s ability to detect the AM sound cue (thereby reducing the hit rate), and/or by increasing indiscriminate responding to the unmodulated noise (thereby increasing the false alarm rate). This latter scenario is important to consider, as OFC lesions have been associated with changes in impulsive behavior (50, 51). To address this issue, we asked how muscimol infusion separately impacted hit rates and false alarm rates. We found that muscimol treatment drastically reduced hit rates in all animals tested (p < 0.001; Figure 3G). In contrast, muscimol treatment only marginally increased false alarm rates (p = 0.039; Figure 3H). This small uptick in false alarm rates might be explained by the fact that animals that receive more frequent punishment are more hesitant to stay on the spout. Indeed, miss rates strongly predicted false alarm rates (p = 0.001; Figure S4B). Collectively, these observations suggest that after OFC inactivation, animals struggled to detect AM noise.

### OFC inactivation prevents task-dependent modulation of auditory cortical neurons

Task-dependent modulations of auditory cortical activity are not fixed in magnitude, and their strength correlates with behavioral AM detection abilities. Strong modulations are associated with excellent AM detection skills, whereas small modulations are associated with poor performance (13). This observation, together with the findings presented above, raises the possibility that OFC inactivation impairs behavioral AM detection by disrupting task-dependent modulations of auditory cortical activity.

To test this hypothesis, we implanted three gerbils (one male) with cannulas in bilateral OFC and a 64-channel silicon probe in left auditory cortex. We initially recorded 22 single- and 45 multi-units in layer 2/3 of the auditory cortex during passive sound exposure. We then infused saline or muscimol into the OFC and recorded from the same auditory cortical units as the animals performed the AM detection task (Figure 4A). Given the relatively small number of single units recorded under each treatment condition (saline: n = 12, muscimol: n = 10), we pooled single- and multi-units together for our statistical analyses. Electrode tracks were visualized offline and used to reconstruct recording site locations (Figure 4B).

**Figure 4.**
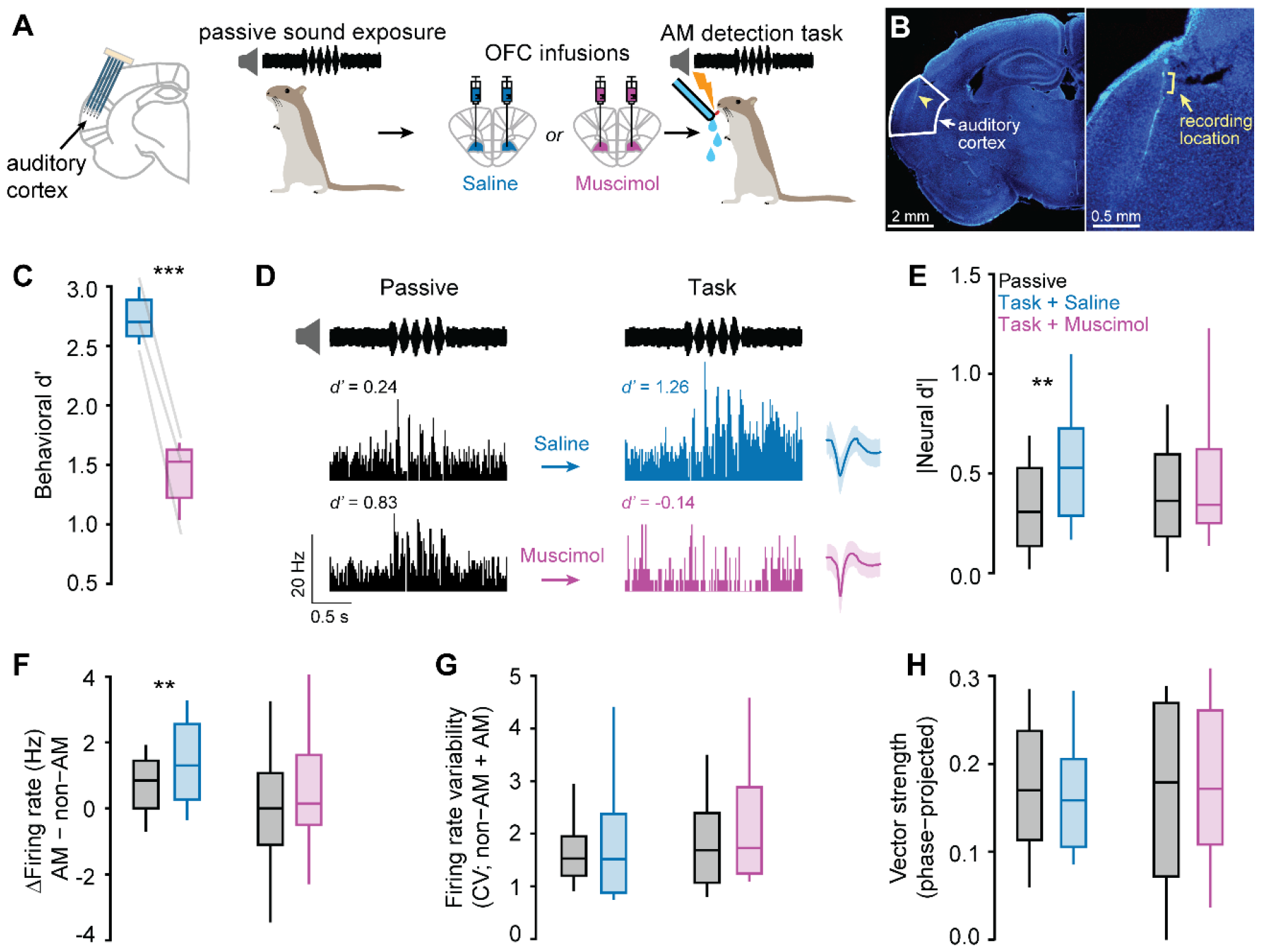
OFC inactivation prevents contextual modulation of auditory cortical neurons. (A) Auditory cortical multi- and single-units were chronically recorded while animals were passively exposed to sounds. Animals then received muscimol or saline infusions in bilateral OFC. Following the infusions, recordings were again made from the same auditory cortical units while animals performed the AM detection task. (B) Electrode placement was confirmed to be in auditory cortex by visualizing fluorescently labeled tracks (green) post-mortem. Recording sites were estimated to be in layers 2/3 (brackets in inset). (C) OFC inactivation impaired behavioral AM detection, replicating the results presented in Figure 3. (D) AM-evoked responses are shown from two representative auditory cortical units recorded from the same animal on different days. Top: An auditory cortical unit recorded before and after saline infusion into the OFC exhibits stronger AM-evoked firing during the task than during passive sound exposure, consistent with our previously published work (13). Bottom: An auditory cortical unit recorded before and after muscimol infusion into the OFC exhibits a modestly reduced firing rate during the task compared to passive sound exposure. (E-F) OFC inactivation prevents a task-dependent increase in auditory cortical neural *d’* values (E) and prevents a task-dependent increase in the separation of AM and non-AM firing rate distributions (F). This separation (termed ΔFiring Rate) was calculated for each unit as the difference between AM-evoked and non-AM-evoked firing rates. (G-H) Neither task engagement nor OFC inactivation affected firing rate variability (G) or phase-locking (H). **p < 0.01, ***p < 0.001.

We first confirmed that OFC inactivation impaired behavioral performance in these animals. As expected from our previous findings, muscimol infusion reduced behavioral *d’* values (p < 0.001; Figure 4C) and hit rates (p = 0.004; Figure S5A) in all subjects without significantly affecting false alarm rates or water consumption (all p > 0.099; Figure S5B-C). We then asked how OFC inactivation affected auditory cortical activity. Figure 4D illustrates AM-evoked responses from two representative auditory cortical units from the same animal recorded on different days. The responses in the top row were recorded from one unit before and after saline infusion into the OFC. This unit showed a stronger AM-evoked firing rate during the task than during passive sound exposure, consistent with our previously published work (13). In contrast, the responses in the bottom row, which were recorded from a different unit before and after muscimol infusion into the OFC, showed a moderately reduced firing rate following OFC inactivation.

To determine whether these differences in response gain translated into differences in AM sensitivity, we transformed the firing rates of individual units into a neural discriminability metric (neural *d’*), using our previously established approach (13, 52). We report absolute neural *d’* values here because negative *d’* values can also support enhanced AM detection via decreases in firing relative to baseline. When saline was infused into OFC, absolute neural *d’* values were significantly higher during task engagement than during passive sound exposure (p = 0.007, Figure 4E), as expected from our prior work (13). In contrast, when muscimol was infused into the OFC, absolute neural *d’* values during task engagement and passive sound exposure were similar (p = 0.162). These data demonstrate that suppressing the OFC prevents task-dependent modulations of the auditory cortex.

The *d’* value for each neuron depends on its average firing rate and its firing rate variability during non-AM and AM noise (see Supplementary Methods in SI Appendix). To determine which of these factors are impacted by OFC inactivation, we first examined the firing rates of individual neurons and compared these values between passive and task sessions. We found that task engagement did not systematically affect auditory cortical firing rates evoked by either non-AM or AM noise, regardless of drug treatment (all p > 0.1; Figure S6A-B). This observation raised the possibility that the differences we observed in neural *d’* values are due to joint changes in non-AM and AM noise responses that manifest on a unit-by-unit basis (see Supplementary Methods in SI Appendix and (13)). To test this idea, we calculated the difference between non-AM and AM firing rates for each auditory cortical unit (here termed the ΔFR), and then compared ΔFR values between passive and task sessions. We found that ΔFR significantly increased during task engagement when saline was infused into the OFC (p = 0.004). In contrast, ΔFR values remained similar between sessions when muscimol was infused (p = 0.121; Figure 4F).

We then calculated the coefficient of variation (CV) for each unit as an indicator of firing rate variability and compared CV values between passive and task sessions. Task engagement did not significantly affect non-AM- or AM-evoked CV values, regardless of drug treatment (all p > 0.4; Figure S6C-D), nor did it affect the sum of non-AM and AM CV values (all p > 0.192; Figure 4G). Together, these findings suggest that OFC inactivation prevents a task-dependent increase in the separation of AM and non-AM firing rate distributions, and not a task-dependent reduction in firing rate variability.

In addition to modulating average firing rates, task engagement can also impact sound encoding and perception by enhancing stimulus phase-locking (27, 43). Therefore, we calculated the phase-projected vector strength evoked by AM noise for individual auditory cortical units and compared these values between passive and task sessions (27, 43, 53). We found that task engagement had no effect on vector strength regardless of drug treatment (all p > 0.625; Figure 4H). This finding is consistent with the observation that task-dependent effects on phase-locking are more significant when AM depths are small (27).

Collectively, these results indicate that OFC inactivation prevents context-dependent enhancements of behaviorally-relevant sound representations in the auditory cortex by modifying neuronal firing rates, without affecting neuronal phase-locking.

### Compensatory mechanisms restore behavioral and auditory cortical AM sensitivity after repeated OFC inactivation

In two subjects (one female) chronically implanted with OFC cannulas and auditory cortex electrodes, we infused saline or muscimol into the OFC multiple times over several days in an attempt to enlarge our sample of auditory cortical neurons. However, we noticed that the effectiveness of repeated muscimol infusions quickly waned and ultimately failed to significantly impact behavior (Figure 5A), indicative of a compensatory mechanism after whole-session OFC inactivation (54, 55).

**Figure 5.**
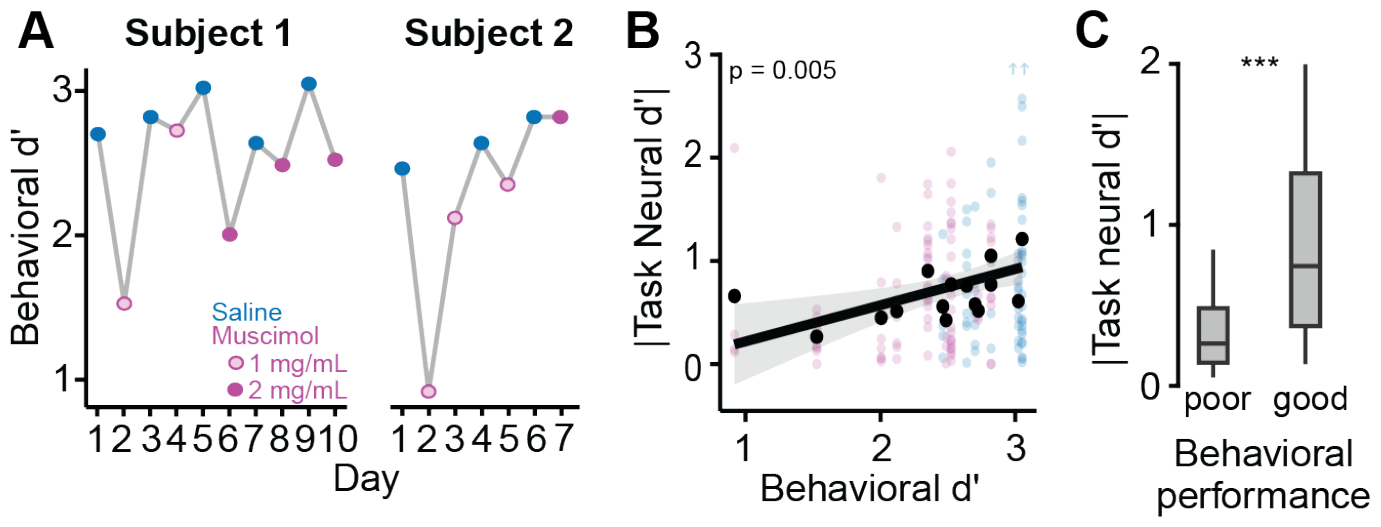
Auditory cortical AM sensitivity correlates with behavioral performance. (A) Repeated muscimol injections into bilateral OFC exert progressively weaker effects on behavior in two subjects. Doubling the muscimol concentration (dark magenta) modestly restored the behavioral effect in one subject (left). (B) Absolute neural d’ values for individual auditory cortical units (small translucent points) are plotted as a function of the behavioral d’ value obtained during the recording session. Black points illustrate median absolute neural *d’* for each behavioral *d’* value. Blue arrows indicate two points that exceeded the depicted range of neural *d’* values. Neural and behavioral d’ values are significantly correlated (see Table 1 for full statistics). (C) Absolute neural *d’* values obtained during the task were significantly higher on days when behavioral performance was good (two best days from each animal) versus days when behavioral performance was poor (two worst days from each animal). n = 30 units on poor performance days and n = 49 units on good performance days. ***p<0.001.

We took advantage of this variability to ask whether muscimol’s effectiveness, inferred from downstream auditory cortical AM sensitivity, predicted behavioral AM detection. We recorded from 60 single- and 96 multi-units over ten days. Indeed, behavioral *d’* significantly correlated with neural *d’* during the task (p = 0.002; Figure 5B). To highlight this relationship, we compared neural *d’* values from the two worst performance days from each animal with the two best days (regardless of drug treatment). We found that neural *d’* values were significantly higher when behavioral performance was good than when performance was poor (p < 0.001; Figure 5C). These findings suggest that OFC is a key part of a distributed network that provides contextual information to the auditory cortex, and when OFC activity is repeatedly suppressed, other brain regions may be recruited to compensate for the loss.

## Discussion

Context-dependent modulation is a widely reported feature of the auditory cortex and is hypothesized to adaptively modify the perceptual salience of stimuli in response to moment-to-moment changes in their behavioral relevance (13, 17, 18, 20, 22, 23, 27, 28, 42–46, 48). Although prior work has identified several brain regions capable of modifying auditory cortical responses and behavior (40, 48, 56–58), our understanding of the neural networks that mediate rapid shifts in auditory perception remains limited. In this study, we reveal that a subset of OFC neurons are sensitive to behavioral context, exhibiting task-dependent dynamics that mirror those previously observed in the auditory cortex (13, 23–25, 30, 49). Our pharmacological experiments demonstrate that OFC inactivation prevents task-dependent enhancements of auditory cortical sensitivity, and, in turn, impairs the ability of animals to detect AM noise. Notably, the behavioral effect of repeated whole-session OFC inactivation gradually waned, suggesting that compensatory brain regions may be recruited when OFC output is unavailable. Together, our results suggest that OFC provides contextual information to the auditory cortex to facilitate the perception of behaviorally relevant sounds.

We found that the OFC is strongly modulated by behavioral context. Both the OFC neuronal population as a whole and the specific subset of neurons that innervate the auditory cortex represent behaviorally relevant cues, such as trial outcome, when gerbils perform a sound detection task. These results are consistent with a recent report from mice (41) and support the idea that the OFC makes an important contribution to auditory cortical sound processing and perception.

During both passive sound exposure and task engagement, OFC neurons were largely unresponsive to AM stimuli. This finding differs from prior work documenting sound-evoked activity in OFC neurons across behavioral states and even under anesthesia (41, 59–61). One possible explanation for this apparent discrepancy is the fact that OFC neurons exhibit relatively poor broadband noise thresholds (e.g. >= ∼50 dB SPL) (60, 61), and all prior studies characterized OFC sound responses at or well above this intensity. Our stimuli, in contrast, were presented at 45 dB SPL and were therefore likely below threshold for most or all OFC neurons. Intriguingly, despite the fact that our acoustic stimuli did not systematically evoke OFC activity, we still observed profound effects of OFC inactivation on auditory cortical plasticity and behavioral output. These results suggest that auditory cortical inputs to the OFC may not be necessary for OFC to successfully signal contextual information back to the auditory cortex. Future experiments could test this hypothesis by silencing auditory cortical axon terminals in the OFC during sound-guided behavior.

We found that OFC inactivation reduces the sensitivity of auditory cortical neurons to behavioral context and impairs behavioral performance of an AM detection task. These data suggest that OFC plays a key role in transmitting contextual information to the auditory cortex to shape sound perception, but the route(s) by which such information is transmitted remains uncertain. Several recent studies support the involvement of a monosynaptic pathway. For example, OFC axon terminals in mouse auditory cortex are sensitive to listening conditions and encode behavioral choice (41). Optogenetic stimulation of these terminals reshapes auditory cortical frequency tuning in anesthetized mice (59), and silencing of these terminals disrupts the ability of animals to flexibly categorize sounds (40). Similar OFC terminal manipulations in the visual or somatosensory cortices alter sensory-evoked responses and impact sensory learning (38, 39). These data suggest that direct projections from the OFC convey contextual information to sensory cortices to modulate stimulus perception.

OFC may also support context-dependent plasticity in sensory cortices via multi-synaptic pathways that recruit neuromodulatory centers. For instance, cholinergic input from the nucleus basalis alters auditory cortical receptive fields and sound perception, and cholinergic axons increase their activity during sound-guided behavior (48, 62–66). Similarly, dopaminergic input from the ventral tegmental area (VTA) influences thalamocortical gain and receptive field organization in the auditory cortex, and is implicated in auditory avoidance learning (67–70). OFC projects to both nucleus basalis (32, 71) and the VTA (72) and regulates VTA firing (73). Thus, OFC neurons may indirectly shape auditory cortical processing and sound perception by relaying contextual information to cholinergic or dopaminergic neurons that innervate the auditory cortex. Targeted manipulations of OFC axons in the nucleus basalis and VTA are needed to resolve these possibilities.

We previously found that task-dependent modulations of auditory cortical neurons grow stronger as animals learn to detect small, near-threshold amplitude modulations (13). This observation suggests that the non-sensory inputs that rapidly modulate auditory cortical activity during task performance also mediate the gradual, longer-term plasticity processes that underlie perceptual learning. This idea is supported by human behavioral experiments demonstrating that in some cases, perceptual learning can be generated by interleaving brief bouts of practice (which are too short to drive learning on their own) with passive sound exposure (74). These data suggest that (I) the non-sensory processes engaged by task performance remain active for some time after task performance ends, and (II) if sounds are passively presented during these periods of residual non-sensory activity, learning can occur. This interpretation closely matches our neurophysiological findings: both auditory cortical (13) and a subset of OFC neurons exhibit task-dependent changes in activity that persist for several minutes after the task has ended. Taken together, these findings support a conceptual framework that task engagement puts sensory cortical neurons into a ‘sensitized’ state permissive for learning, which lasts for many minutes beyond the end of the task (13, 23, 75, 76). The neural circuits involved in regulating this phenomenon are still unclear, but our data suggest that a subpopulation of OFC neurons may be involved. Future work should use viral approaches to identify the downstream targets of these neurons, and to determine their causal contributions to task-induced sensitization of sensory circuits and practice-based improvements in auditory perception.

## Materials and Methods

For a full description of the methodology, see Supplementary Materials in the SI Appendix.

### Subjects

Adult Mongolian gerbils (*Meriones unguiculatus*) used in this study were bred in-house from commercially obtained breeding pairs (Charles River). Animals were kept under 12 hr light:12 hr dark cycle and on *ad libitum* food, enrichment and water access until placed on controlled water access for behavioral experiments. All procedures were approved by the Institutional Animal Care and Use Committee at the University of Maryland College Park.

### Behavioral task and passive sound exposure

Auditory perception was tested with an aversive go/no-go amplitude-modulated (AM) detection task as previously described (13, 52, 77–80). Briefly, unmodulated broadband noise was continuously presented from a calibrated speaker and unpredictably transitioned into AM noise while animals drank from a water spout. AM detection performance was assessed by *d*’ = *z*(*HR*) − *z*(*FAR*), where z(HR) and z(FAR) are the z-scores at the probability levels of hit rates (HR) and false alarm rates (FAR). For passive sound exposure, the waterspout was removed from arena, and unmodulated noise transitioned to AM noise pseudorandomly every 3-5 s.

### Electrophysiology

Extracellular neural activity was recorded chronically from either left OFC or left auditory cortex. After spike sorting, auditory cortical firing rates were transformed to neural *d’* (13, 46). The ability of the units to phase-lock to the 5 Hz AM stimulus was analyzed by calculating the phase-projected vector strength (53). The backs of electrode shanks were painted with azide-free fluorescent microspheres (FluoSpheres 0.2 μm; ThermoFisher) before implantation to verify implant site location *post mortem*.

### Software and analysis

Statistical analyses were performed using R (version 4.3.1) and RStudio (version 2023.9.1.494; Posit Software). All statistical results can be found in Table 1. Boxplots center lines represent medians, boxes represent 25-75% interquartile range and whiskers represent 10-90% interquartile ranges.

### Brain infusions

Muscimol (1-2 μg/μL; 0.25-0.5 μL/hemisphere) or vehicle (0.9% saline) was infused through chronic bilateral cannula targeting OFC ∼40 min before the task.

## Acknowledgments

We thank current and former members of the Caras Lab for helpful feedback and discussions and assistance with animal care. Special thanks to Dr. Daniel Stolzberg for assistance with software for behavioral testing. We acknowledge the Imaging Core Facility in the department of Cell Biology and Molecular Genetics at the University of Maryland, College Park for training and the use of the Zeiss LSM 980 Airyscan 2 confocal microscope. Purchase of the Zeiss LSM 980 Airyscan 2 was supported by Award Number 1S10OD025223-01A1 from the National Institute of Health (NIH). This work was supported by NIH R00DC016046 and R01DC020742 to MLC.

## Supporting Information Text

## Supplementary Methods

### Behavioral set-up

Auditory perception was tested with an aversive go/no*-*go AM detection task as previously described (1–4). A custom test cage (CCMI Plastics) containing a stainless-steel water spout and metal floor plate was positioned inside a sound attenuating booth (GretchKen). Infrared detection of the animal at the spout triggered water delivery via a programmable syringe pump (NE-1000, New Era Pump Systems). Sound stimuli were delivered via a calibrated free-field speaker (DX25TG59-04 1” Fabric Dome Tweeter, Peerless by Tymphany) positioned directly above the test cage. Shocks were delivered by an H13-15 Precision Animal Shocker (Colburn). Behavior was monitored remotely via webcam. Data acquisition, sound delivery, and digital outputs were controlled via an RZ6 multifunction processor (Tucker Davis Technologies, TDT).

### Behavioral training and testing

During an initial procedural training stage, animals were placed on controlled water access and learned to drink steadily from a spout while in the presence of a “safe cue” (unmodulated broadband noise, 0.1-20 kHz, 45 dB SPL). Animals were then trained to withdraw from the spout when the sound changed to the “warn” cue (5 Hz sinusoidal amplitude modulated (AM) noise, 0 dB re: 100% depth, 1 second duration) by pairing the warn cue with a mild shock (0.5-1 mA, 300 msec). Warn trials were randomly interspersed with 3-5 safe trials, but only while animals made contact with the spout. The gain of the AM signal was adjusted to control for changes in average power (5).

The animal’s contact with the spout was monitored during the final 100 msec of each trial. Breaking spout contact for at least 50 msec during the monitoring window was considered a spout ‘withdrawal,’ and was scored as a ‘hit’ on AM trials and as a ‘false alarm’ on non-AM trials. Behavioral performance was assessed by *d*^′^ = *z*(*HR*) − *z*(*FAR*), where z(HR) and z(FAR) are the z-scores of the hit rates (HR) and false alarm rates (FAR), respectively. To avoid values that approach infinity, extreme HR and FAR values were set to a floor of 5% and a ceiling 95%. Behavioral testing commenced only after an animal achieved a *d’* >= 2 in a single session.

For passive sound exposure sessions, animals were placed in the test cage but did not have access to a waterspout. All other elements of the experimental apparatus were identical to the detection task. Sounds were presented in a manner similar to that of the task (AM stimuli randomly interspersed with 3-5 seconds of unmodulated noise).

Note that the fourteen animals used for the OFC electrophysiology and fiber photometry experiments (Figures 2 and S1-3) were recorded from as they underwent a perceptual learning protocol in which they were trained to detect progressively smaller AM depths. The details of this procedure have been previously described (6). OFC’s involvement in perceptual learning will be explored in a future manuscript.

### Electrode implant procedures

NeuroNexus 64-channel probes (OFC: A4x16-Poly2-5mm-20s-lin-160; auditory cortex: Buzsaki64_5x12) were mounted on custom-made microdrives. The backs of the electrode shanks were painted with azide-free fluorescent microspheres (FluoSpheres 0.2 μm; ThermoFisher) just before implantation. On the day before surgery, animals were placed on antibiotic treatment (minocycline in drinking water; 0.02 mg/mL) and given meloxicam (1.5 mg/kg). On the day of the surgery, they were given another dose of meloxicam plus dexamethasone (0.35 mg/kg). Then, they were anesthetized with isoflurane (1-5%) and secured in a stereotaxic apparatus (Kopf). Ophthalmic ointment was applied to the eyes and alternating swabs of alcohol and betadine were applied to the skin on the top of the head. The skull was exposed and dried with hydrogen peroxide, three or four bone screws were inserted, and craniotomy locations were marked over left OFC or left auditory cortex after leveling the head. Note that we leveled the animal’s head between lambda and bregma, rather than between lambda and the occipital ridge as it was done for the gerbil brain atlas (7). Therefore, all of our rostro-caudal coordinates differ from those reported in the atlas.

Before auditory cortex craniotomy, the temporalis muscle insertion was detached, and muscle was separated from skull using dry SurgiFoam (Ethicon). The skull and screws were covered with C&B Metabond (Parkell) for better implant adhesion. The edges of the craniotomy were thinned by drilling and the bone flap was removed. A durotomy was made over the area of interest and electrodes were implanted using the following coordinates (all relative to lambda). OFC: 1.20–1.50 mm lateral, 9.05–9.50 mm rostral, 2.00-2.50 mm ventral, angle=0° from vertical; implanted along coronal plane. Auditory cortex: 4.80 mm lateral, 3.90 mm rostral (anterior shank), 1.10-1.50 mm ventral, angle=20° clockwise from vertical plane facing the animal’s posterior side, implanted along parasagittal plane. A ground wire was inserted into the right caudal hemisphere. Implants were secured with dental cement (Palacos). The day following surgery, animals were given another dose of meloxicam and dexamethasone. Antibiotic treatment continued for an additional six days. Controlled water access and behavioral experiments started at least seven days after surgery.

### Electrophysiology

Extracellular recordings from the OFC were obtained from freely-moving animals before (passive-pre), during (task), and after (passive-post) behavioral testing sessions that took place every 24-48 hours. The number of AM stimulus presentations was similar across behavioral contexts (passive-pre:100 ± 0.10; task: 110 ± 3.27; passive-post: 99.8 ± 0.23 trials; means ± SEMs).

Extracellular recordings from the auditory cortex were obtained before (passive-pre) and during (task) behavioral testing sessions that took place every 24-48 hours. The number of AM stimulus presentations was similar across behavioral contexts (passive-pre: 38.3 ± 8.08; task: 42.5 ±7.05; means ± SEMs).

Neurophysiological signals were obtained via one of two methods: (I) Analog signals were acquired with a 64-channel wireless headstage and receiver (W64, Triangle BioSystems), preamplified and digitized at 24.4 kHz sampling rate (PZ5; Tucker Davis Technologies, TDT), and integrated with behavioral timestamps via RZ2 and RZ6 processors controlled by the Synapse software suite (TDT). Raw signals were sent to an RS4 data streamer (TDT) and stored online. (II) Signals were preamplified and digitized at 30 kHz by a 64-channel RHD headstage (Intan Technologies) connected to a slip-ring commutator (Taidacent; purchased from Amazon) and integrated with behavioral timestamps using an RHD recording controller (Intan) and OpenEphys software (8).

Offline signals were high-pass filtered at 150 Hz and common-median referenced across channels. Multi- and single-unit sorting was performed using Kilosort2 (9). Sorting results were manually curated using Phy (https://github.com/cortex-lab/phy). Clusters were classified as putative single-units during manual curation when they presented with a high signal-to-noise ratio, clear refractory period gap in the autocorrelogram and clear cross-correlogram isolation from neighboring clusters. After manual curation, the Allen Brain Institute quality metrics ‘presence ratio’, ‘amplitude cutoff fraction missing’ and ‘ISI violation false-positive rate’ (parameters: isi_threshold = 1.5 ms; min_ISI = 0.15 ms) (10) were calculated (https://allensdk.readthedocs.io/en/latest/_static/examples/nb/ecephys_quality_metrics.html). We also calculated the ‘refractory period violation rate’ as the percentage of interspike intervals that occurred within the refractory period of 1.5 msec. Only putative single-units that had a ‘presence ratio’ > 0.9, ‘amplitude cutoff fraction missing’ < 0.1, ‘ISI violation false-positive rate’ < 0.5, and ‘refractory period violation rate’ < 2% were considered true single-units.

After single- and multi-unit isolation, firing rates were calculated. The window used for calculating firing rates on “Hit” trials was the full 1-s AM duration. “Miss” trials produced a shock artifact, however. Thus, the window used for calculating firing rates on “Miss” trials was the first 950 msec of the AM stimulus. “False alarm” trials by definition do not contain an AM period, so we used a 1-s block of non-AM sound during which a false alarm occurred to quantify firing rates. Peristimulus time-histograms (PSTH) were generated using 10-20 msec bins.

Auditory cortical firing rates were transformed to neural *d’* (6, 11) using the formula:

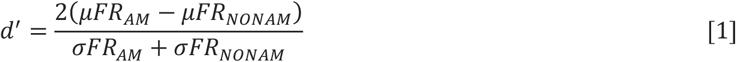

where *µFR*_*AM*_ and *µFR*_*NONAM*_ are the mean firing rates and *σFR*_*AM*_ and *σFR*_*NONAM*_ are the standard deviation of the firing rates during AM and non-AM presentations, respectively. The non-AM period used for these calculations was generally the 1 s non-AM noise period immediately preceding an AM trial. However, if animals withdrew from spout during that period (i.e., false alarm), the nearest previous 1 s period during which the animal did not leave the spout (i.e., correct rejection) was used.

The effect of task engagement on OFC firing rates during non-AM sound presentation was assessed with a Wilcoxon rank-sum test. When p < 0.05, the W statistic was used to determine the direction of the change: ‘task-enhanced’ units exhibited a W > 0, while ‘task-suppressed’ units exhibited W < 0. The magnitude of the context modulation was computed for each OFC unit using the z-score formula:

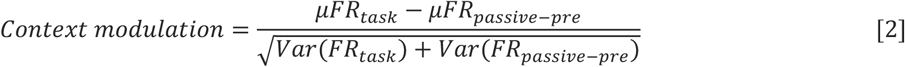

where *µFR*_*task*_ and *µFR*_*passive-pre*_ are the mean firing rates and *Var*(*FR*_*task*_) and *Var*(*FR*_*passive-pre*_) are the firing rate variances during the task and during the passive-pre periods, respectively.

The response of OFC neurons to AM sound presentation was assessed by calculating the area under the receiver operating characteristic curve (auROC) from peri-stimulus time histograms (PSTH; Figure S1A-B), as previously described (12). This transformation is useful for comparing response dynamics across many units with widely different firing rates. Briefly, PSTHs were generated using 10-ms bins starting 2 s before until 5 s after trial onset (Figure S1A). We compared the histogram of firing rates during a baseline period (from 2 to 1 s before onset) to the histogram during 100-ms windows along the entire PSTH (Figure S1C-D). For these comparisons, a moving criterion vector 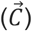 was created ranging from 0 to maximum firing rate in the PSTH in 100-ms increments. For each value *c* in 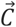, we calculated the probability that the activity during each bin was higher than *c* and plotted it against the probability that the activity during baseline was higher than *c*, constructing an ROC curve (Figure S1E-F). The auROC was calculated via trapezoidal numerical integration. AuROC values range from 0 to 1, and values above 0.5 indicate relative increases, while values below 0.5 indicate relative decreases in firing.

### Vector strength

Phase-projected vector strength was calculated as in (13). Briefly, the standard trial-by-trial vector strength (*VS*_*t*_) was calculated as

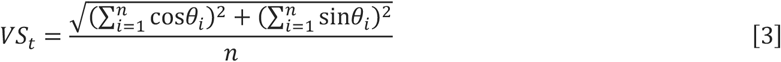

where n is the number of spikes in a given trial and *θ*_*p*_ is the phase of each spike in relation to the stimulus modulation period. *θ*_*i*_ was calculated as

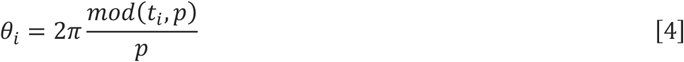

where *mod*(*t*_*i*_,*p*) is the modulo (division remainder) between the spike time relative to the onset of the stimulus *t*_*i*_ and the modulation period for a 5 Hz stimulus *p* (i.e., 0.2 s). The phase-projected vector strength (*VS*_*pp*_) was calculated as

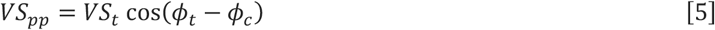

where *ϕ*_*t*_ and *ϕ*_*c*_ are the trial-by-trial and mean phase angle, calculated as

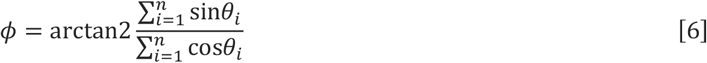

where n is the number of spikes in each trial (*ϕ*_*t*_) or in all trials (*ϕ*_*c*_), and arctan2(y/x) yields the angle between the x-axis and the vector from [0, 0] to [x, y] (base R function). *VS*_*pp*_ ranges from –1 (all spikes in each trial are out of phase with the mean response) to 1 (all spikes are in phase with the mean response).

### Fiber photometry

Pre-operative care was as described above. The skull was exposed, dried with hydrogen peroxide, and leveled between lambda and bregma. Two bone screws were inserted, and craniotomy locations were marked over left OFC and left auditory cortex. The temporalis muscle was detached as described above. A virus (AAVrg-syn-jGCaMP8s-WPRE; Addgene; (14)) was injected into the left auditory cortex to drive retrograde expression of a genetically-encoded calcium indicator (jGCaMP8s). Injections were made at three locations along the rostro-caudal axis between 2.40 and 3.90 mm rostral to lambda. At each rostro-caudal location, injections were made at two depths relative to the pial surface (0.90 and 0.30 mm; 200 nL per depth), totaling 1200 nL. Each injection was made directly into the temporal aspect of the brain, with the pipette angled 45 degrees laterally. The auditory cortex craniotomy was covered with KwikSil (WPI) and the skin was repositioned to cover the craniotomy. Skin margins were glued with cyanoacrylate tissue adhesive and bone screws and skull were covered with Metabond. Then, a craniotomy over the left OFC was made and an optical fiber (400 μm core; ø1.25 mm ceramic ferrule; 0.5 NA; RWD Life Science) was implanted in the left OFC using the following coordinates (relative to lambda): 1.2 mm lateral, 8.90-9.05 mm rostral, 2.90-3.20 mm ventral from pial surface). The craniotomy was covered with KwikSil and the fiber was secured with dental cement. Post-operative care was as described above. One control animal (female) was injected with AAVrg-hSyn-eGFP (Addgene) in the left auditory cortex and had a fiber implanted in the left OFC as described above. Controlled water access and behavioral experiments started at least three weeks after surgery to allow for virus expression. Fiber photometry recordings were performed four weeks after surgery.

All optical equipment (patch cords, fluorescence MiniCube, 405 and 465 nm connectorized LEDs, LED driver and Newport photoreceiver module) were purchased from Doric Lenses. LED driver control and photovoltaic signal recording was sampled at 1 kHz using an RZ2 processor and Synapse software (Tucker-Davis Technologies).

Fluorescence was recorded using a 465 nm LED. Calcium-independent (i.e., isosbestic) signals were also recorded using a 405 nm LED to correct for movement artifacts and fluorescence decay. The isosbestic point of GCaMP is ∼410 nm (15).

Before each recording day, patch cords were photobleached overnight. Animal fiber-optic cannulas were connected to the system using a quick-release connector (ADAL3; ThorLabs). Light intensity at the tip of the animal-connecting patch cord was measured with a photodiode power sensor and energy meter (S120C and PM100USB; ThorLabs). Light output (465 and 405 nm respectively) was ∼90 and 28 μW for all animals except for one male (∼45 and 28 μW).

Photometry signals were processed offline similarly to (16). Briefly, 465 and 405 nm signals were 5 Hz low-pass filtered and smoothed with a 100-point zero-phase moving average filter. Then, LED onset artifacts were removed, baseline fluctuations were corrected with the adaptive iteratively reweighted Penalized Least Squares algorithm (airPLS) (17) and signals were standardized by median subtraction and division by standard deviation. The standardized 405 signal was fitted to the standardized 465 signal with non-negative robust linear regression and the fit was subtracted from the standardized 465 signal, resulting in a dF/F signal. Finally, this resulting dF/F signal was aligned to behaviorally relevant timestamps and z-scored using a baseline period ranging from 1 s before stimulus until stimulus onset.

### Brain cannulation and infusions

Pre- and post-surgical care were as described above. The skull was exposed and dried, and two bone screws were inserted. The skull and screws were covered with C&B Metabond (Parkell) for better implant adhesion. Craniotomies and durotomies were performed over OFC. Double guide cannulas (26-gauge, 5 mm length, 1.5 mm center-to-center distance; PlasticsOne model C235GS-5-1.2/SPC) were inserted just above OFC using the following coordinates (all relative to lambda): lateral = 1.50 mm, rostral = 8.75-9.05 mm, ventral = 1.50 mm; angle=0° from vertical) and secured with dental cement (Palacos). Dummy cannula (33-gauge, 0.25 mm protruding from guide cannula; PlasticsOne model C235DCS-5/SPC) were inserted into the guides and secured in place with a dust cap.

Starting at least seven days after surgery, animals were placed on controlled water access and trained to detect AM sounds as described above. Intracortical infusions only began after animals reached expert performance on the task (d’≥2). Muscimol (abcam) was dissolved in in 0.9% saline to achieve a concentration of 1-2 μg/μL. Infusion cannulas (33 gauge, protruding 1 mm from guide cannula; PlasticsOne model C235IS-5/SPC) containing muscimol or saline were inserted into the guides. Bilateral OFC infusions (0.25-0.5 μL/hemisphere, 0.2 μL/min rate) were made simultaneously via a six-channel programmable pump (NE-1600, New Era) and 10 μL glass syringes (Hamilton 1801). Infusions were performed ∼40 min before the behavioral task under light isoflurane anesthesia. The entire infusion procedure lasted between 6 and 10 minutes and animals typically recovered from anesthesia within 2 minutes.

To estimate drug spread and cannula placement, fluororuby (Dextran, Tetramethylrhodamine, 10,000 MW, Lysine Fixable, Invitrogen) was diluted to 5% in sterile saline and infused through the cannulas under light isoflurane anesthesia at the end of the study. Immediately after infusions animals were perfused and brains were processed for histology as described below.

### Histology and imaging

At the end of all experiments, animals were deeply anesthetized with a mixture of ketamine (150 mg/kg) and xylazine (6 mg/kg) in sterile saline and transcardially perfused first with ∼20 mL of phosphate buffered saline (PBS) then with ∼20 mL of 4% paraformaldehyde (PFA) in phosphate buffered saline. After perfusion, implants were removed, and brains were extracted and postfixed in PFA for 1-7 days until sectioning. Brains were then transferred to PBS, embedded in 6% agar, and sectioned on a vibratome (Leica VT-1000S) at 70 μm thickness. Slices were collected and mounted on gelatinized slides and coverslipped with ProLong Gold or Diamond mountant (Molecular Probes).

GFP-containing tissue (photometry experiment) was processed for GFP immunofluorescence for signal amplification. First, free-floating sections were washed in PBS 3x10 min. Non-specific binding was blocked by incubation in 10% normal goat serum (NGS) in phosphate buffered saline containing Triton-X (Sigma; 0.3% PBT) for 2 h at room temperature. Following the blocking step, slices were incubated in 1:1000 rabbit anti-GFP primary antibody (ThermoFisher #A11122) diluted in the same blocking solution (NGS + 0.3% PBT) for 1 h at room temperature then 40-48 h at 4° C. Following primary incubation, slices were washed in 0.1% PBT 3x15 min then incubated in 1:1000 goat anti-rabbit Alexa-488 secondary antibody (ThermoFisher #A32731) for 1 h at room temperature. Finally, following secondary incubation, slices were washed in 0.1% PBT 3x10 min and coverslipped with ProLong Diamond.

Sections were imaged either via epifluorescence (Leica DM750) or confocal microscopy (Zeiss LSM 980 Airyscan 2) at 10x magnification. For epifluorescence images, tiles were taken manually and stitched together using Photoshop 2023 panorama PhotoMerge function (Adobe).

### Software and analysis

We used the ePsych software developed by Dr. Daniel Stolzberg (https://github.com/dstolz/epsych) for MatLab R2014a (MathWorks) for behavioral data acquisition and control.

All data were processed offline by custom MatLab pipelines (https://github.com/caraslab/caraslab-behavior-analysis, https://github.com/caraslab/caraslab-spikesortingKS2, and https://github.com/caraslab/caraslab-fiberphotometry), then further analyzed by custom Python and R routines (available upon request).

Statistical analyses were performed by generalized linear models followed by ANOVA (GLM/ANOVA) using the ‘glmmTMB’ (18) and the ‘car’ (19) packages for R. Normality of GLM residuals was assessed after each fit by visually inspecting the distribution of studentized residuals (q-q plots) using the ‘DHARMa’ package for R (20). When normality was not fulfilled, data were square-root- or log-transformed, and GLMs were rerun. Before transformation, data containing negative numbers were rescaled using the formula 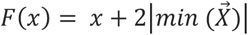, where x represents an individual value and 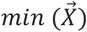 represents the lowest value in the dataset. When interactions were significant, Bonferroni multiple comparison post-hoc tests using the package ‘emmeans’ for R (https://github.com/rvlenth/emmeans) were used after “regridding” the data into original units if a transformation was required.

All GLM fixed effects can be found in the statistical tables. GLM random effects were as follows. The effect of behavioral context on OFC activity included Unit nested under Subject (Figure 2F) or Context (passive-pre, task, passive-post) nested under Unit nested under Subject as random effects (Figure 2G and S3C). The effect of OFC suppression on behavioral AM detection included Subject as random effect (Figures 3C-H, 4C and S4-5). The effect of behavioral context on auditory cortical activity was separately analyzed for each drug treatment (saline or muscimol) including Context (passive, task) nested under Unit nested under Subject as random effects (Figure 4E-H and S6), or Unit nested under either Behavioral *d’* (Figure 5B) or performance (good vs poor, Figure 5C) nested under Subject as random effects. All supplementary statistical results can be found in Table S1.

## Supplementary figures and tables

**Figure S1.**
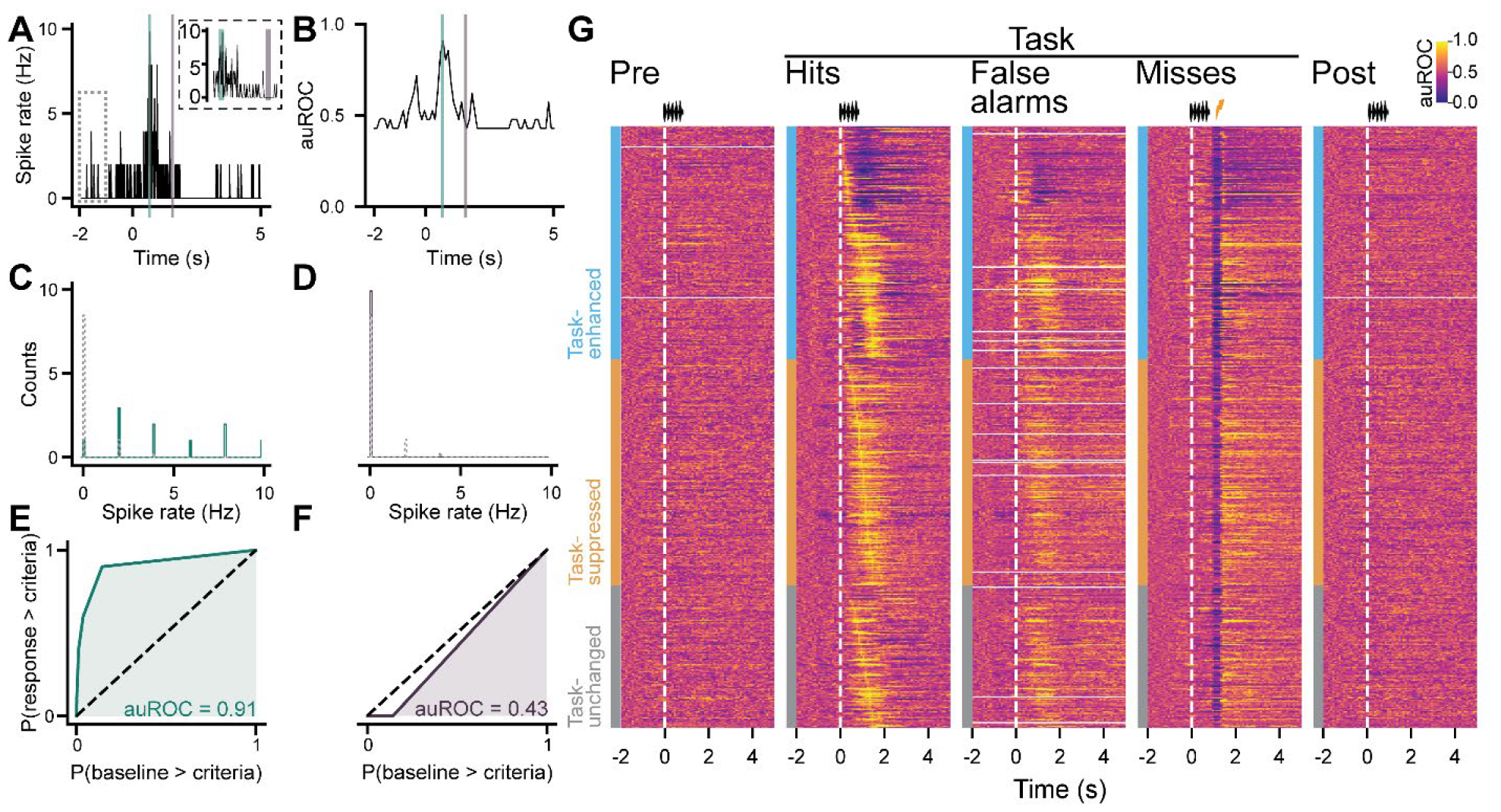
OFC neurons exhibit heterogeneous firing rate changes during sound-guided behavior. (A-B) Peri-stimulus time histograms (PSTH) around trials (A) were used to generate normalized firing auROC PSTHs (B). We compared the spike rate during a baseline period (dashed gray box) to the spike rate during 100-ms windows along the entire PSTH. Two example windows are shown in green and purple. Inset in A represents a magnified view around the windows to illustrate that each 100-ms window contains 10x 10-ms spike rate values. (C-D) Count histograms illustrating the spike rate bins during baseline period (dashed lines) and during the 100-ms example windows (green and purple solid lines) from the PSTH illustrated in A. Spike rate distributions were compared using a criterion vector (see Supplementary Methods for details) constructing ROCs (E-F), from which auROCs could be calculated. (G) Heatmaps illustrate normalized neuronal firing (auROC) aligned to trial onset (Time = 0 s; vertical dashed white lines). Units (rows) are separated by the type of modulation exhibited during non-AM sound presentation (task-enhanced, -suppressed or -unchanged as in Figure 2). Rows are ordered by time of the maximum absolute auROC change since AM trial onset during Hit trials, and the same order is used across other trial types. False alarm trials by definition do not contain an AM period, so Time = 0 s represents the onset of a 1-s trial block within which false alarms occurred. Data are from 298 single- and 227 multi-units. Solid white rows are from sessions during which auROC could not be calculated due to spikes not occurring that period.

**Figure S2.**
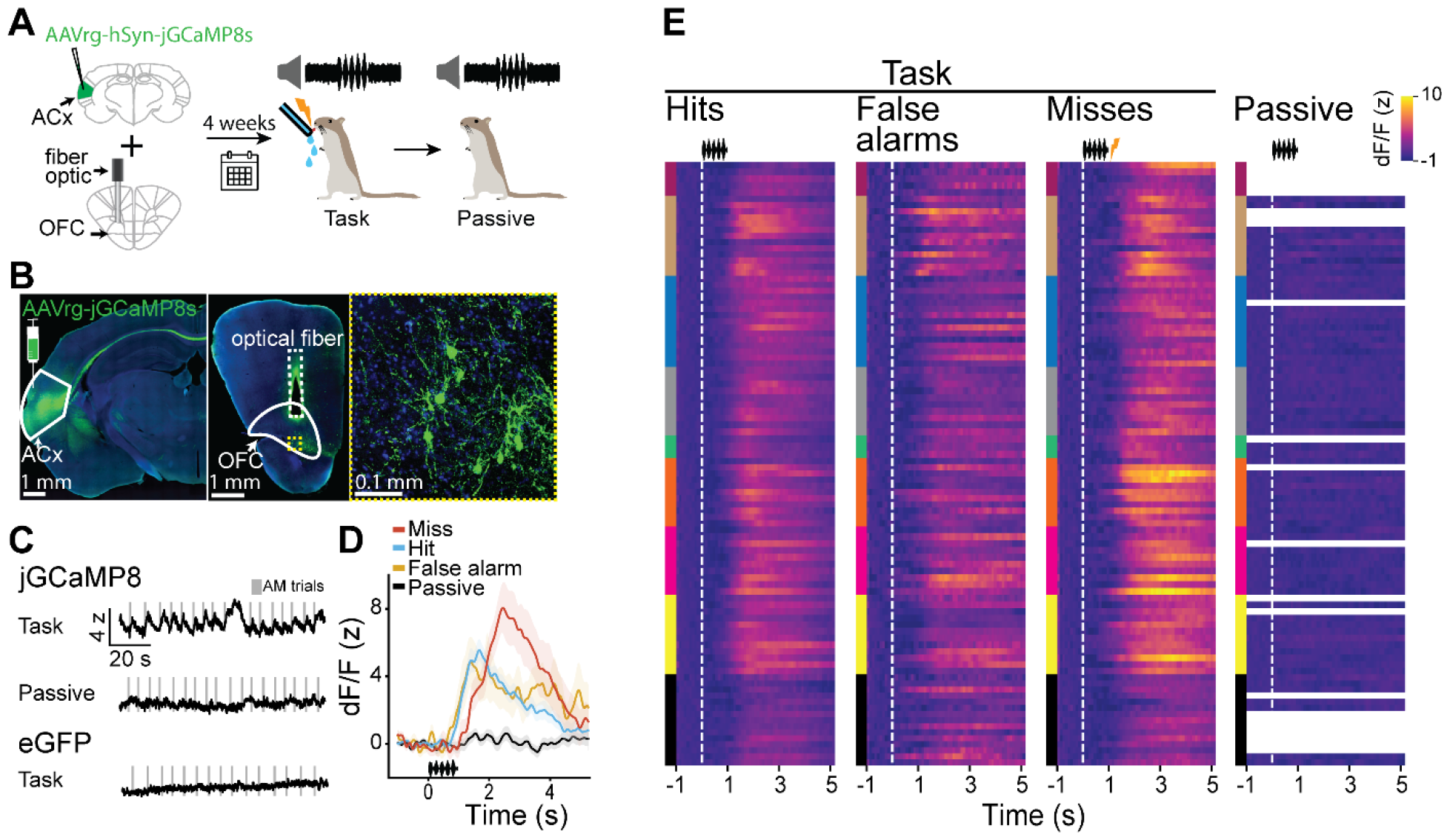
OFC neurons that innervate auditory cortex show increased trial-based activity during task performance. (A) Calcium transients from OFC neurons that project to auditory were recorded via fiber photometry in nine animals (four females). Auditory cortex was injected with a virus retrogradely-expressing jGCaMP8s and an optical fiber was implanted in left OFC. After four weeks to allow for virus expression, animals were placed on controlled water access and tested on the AM detection task. Photometry signals were collected during the task and during a period of passive sound exposure immediately following the task. (B) Virus injection site in auditory cortex (left) and fiber location in OFC (middle and right) were confirmed after the experiment. (C) Calcium transients time-locked to AM trials were only evident during task engagement (top) and not during passive sound exposure (middle). To confirm that the recorded signals were not related to movement artifacts, one female was injected with a virus retrogradely-expressing enhanced green fluorescent protein (eGFP) and a fiber optic was placed in OFC. No task-related transients were detected in this animal (bottom). (D) Representative mean ± standard error calcium transients from one recording session demonstrating calcium dynamics around behavioral responses and not during passive AM presentation. (E) Heatmaps illustrate calcium dynamics around behavioral responses and AM presentations during task and passive periods. Responses were aligned to trial onset (Time = 0 s; vertical dashed white lines). False alarm trials by definition do not contain an AM period, so Time = 0 s represents the onset of a 1-s trial block within which false alarms occurred. Each row represents one session. Vertical color bars represent sessions from the same animal. Since calcium transients were not evident during passive sound playback, recordings were not performed for every session. Blank rows in the passive session columns represent sessions in which no passive recording was made.

**Figure S3.**
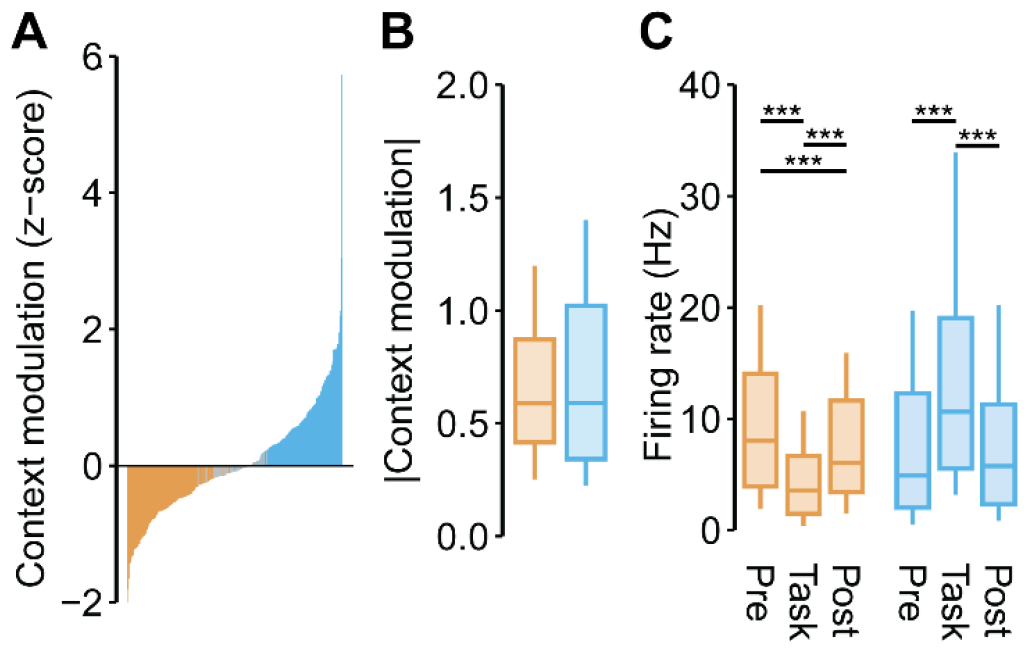
Context-dependent dynamics in non-AM firing are present even when the analysis is restricted to single-units. Same analysis as in Figure 2E-G but restricted to single-units (n = 117 task-enhanced and n = 116 task-suppressed out of 298 single-units). (A) Context modulation values are presented for all single-units sorted in increasing order. Gray bars represent units without significant context modulation. (B) Absolute modulation magnitudes were similar between task-enhanced and - suppressed single-units. (C) Task-suppressed single-units exhibited reductions in tonic activity that persisted in the passive-post period. The activity of task-enhanced single-units, on the other hand, returned to passive-pre levels after the task ended. ***p<0.001

**Figure S4.**
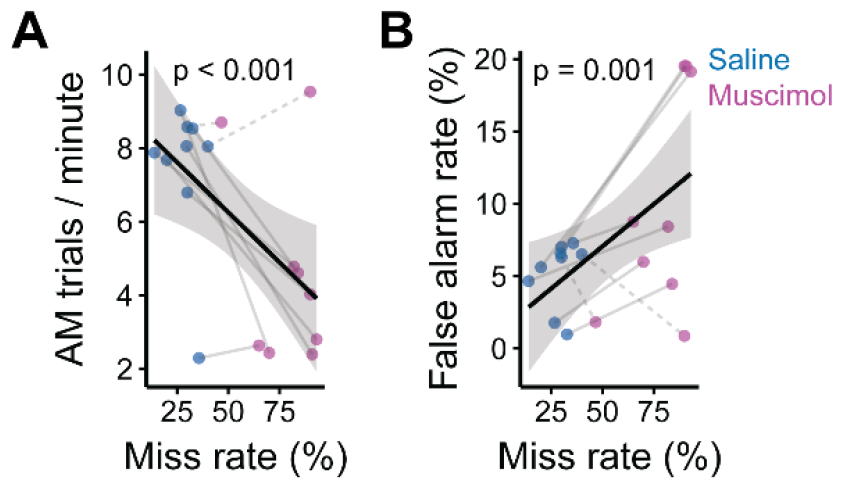
Slower trial completion and increased false alarm rates during OFC inactivation may be explained by the increased frequency of punishment. (A) The number of AM trials completed per minute negatively correlates with the miss rate. (B) The false alarm rate positively correlates with the miss rate. Blue dots represent days when animals received saline infusions into OFC, while magenta dots represent days when animals received muscimol infusions. Gray lines connect data from the same animal. Dashed lines indicate animals that received unilateral left OFC infusions due to partially clogged cannulas.

**Figure S5.**
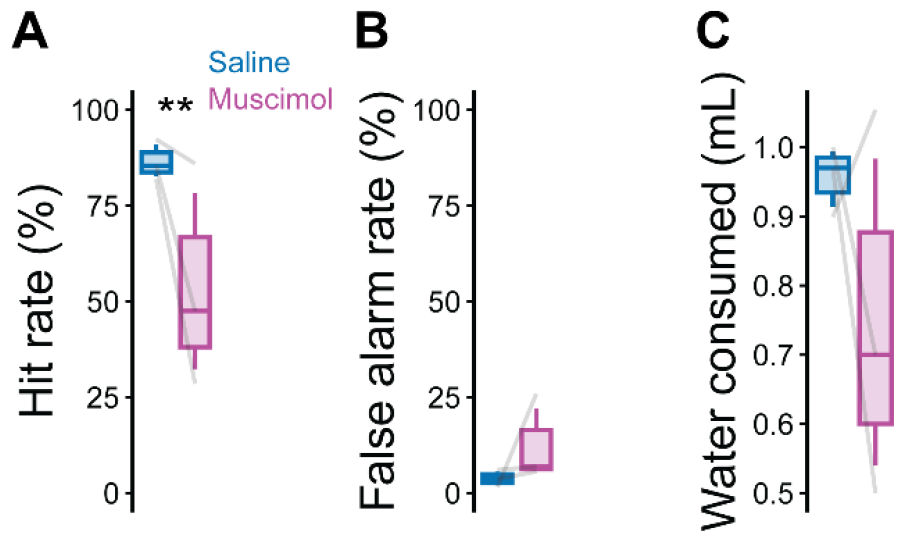
The behavioral effects of OFC suppression replicate in animals with chronic electrode implants in the auditory cortex. OFC suppression significantly reduced behavioral hit rates (A) but did not significantly affect false alarm rates (B) or water consumed during the task (C). **p<0.01.

**Figure S6.**
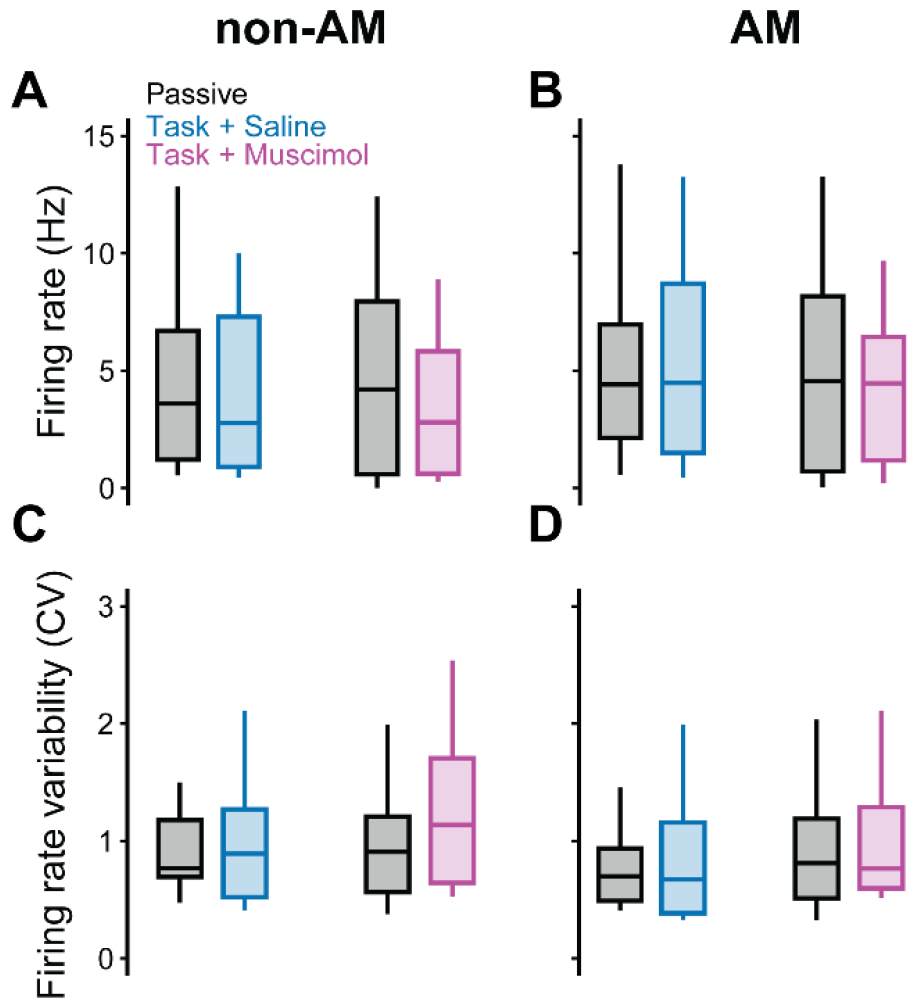
Effects of OFC suppression on auditory cortical firing rates and firing rate variability. (A-B) Neither task engagement nor OFC suppression systematically affect non-AM (A) or AM firing rates (B) of auditory cortical single- and multi-units. (C-D) Neither task engagement nor OFC suppression systematically affect non-AM (C) or AM firing rate variability (D).

**Table S1.**
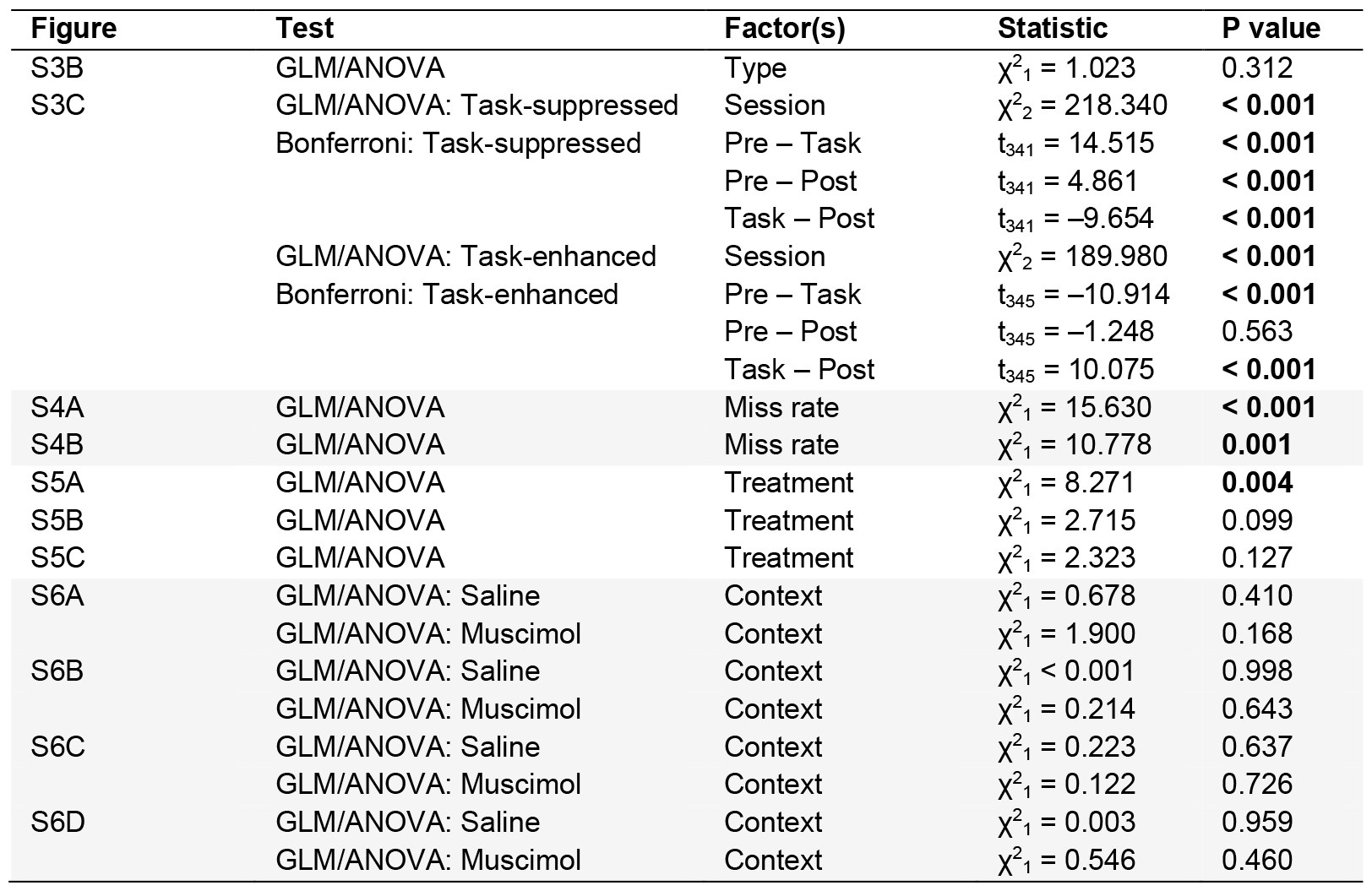
Statistical results.

